# A multi-scale coevolutionary approach to predict interactions between protein domains

**DOI:** 10.1101/558379

**Authors:** Giancarlo Croce, Thomas Gueudré, Maria Virginia Ruiz Cuevas, Victoria Keidel, Matteo Figliuzzi, Hendrik Szurmant, Martin Weigt

## Abstract

Interacting proteins and protein domains coevolve on multiple scales, from their correlated presence across species, to correlations in amino-acid usage. Genomic databases provide rapidly growing data for variability in genomic protein content and in protein sequences, calling for computational predictions of unknown interactions. We first introduce the concept of *direct phyletic couplings*, based on global statistical models of phylogenetic profiles. They strongly increase the accuracy of predicting pairs of related protein domains beyond simpler correlation-based approaches like phylogenetic profiling (80% vs. 30-50% positives out of the 1000 highest-scoring pairs). Combined with the direct coupling analysis of inter-protein residue-residue coevolution, we provide multi-scale evidence for direct but unknown interaction between protein families. An in-depth discussion shows these to be biologically sensible and directly experimentally testable. Negative phyletic couplings highlight alternative solutions for the same functionality, including documented cases of convergent evolution. Thereby our work proves the strong potential of global statistical modeling approaches to genome-wide coevolutionary analysis, far beyond the established use for individual protein complexes and domain-domain interactions.

**Author summary:** Interactions between proteins and their domains are at the basis of almost all biological processes. To complement labor intensive and error-prone experimental approaches to the genome-scale characterization of such interactions, we propose a computational approach based upon rapidly growing protein-sequence databases. To maintain interaction in the course of evolution, proteins and their domains are required to coevolve: evolutionary changes in the interaction partners appear correlated across several scales, from correlated presence-absence patterns of proteins across species, up to correlations in the amino-acid usage. Our approach combines these different scales within a common mathematical-statistical inference framework, which is inspired by the so-called direct coupling analysis. It is able to predict currently unknown, but biologically sensible interaction, and to identify cases of convergent evolution leading to alternative solutions for a common biological task. Thereby our work illustrates the potential of global statistical inference for the genome-scale coevolutionary analysis of interacting proteins and protein domains.

## Introduction

Essential to life at the molecular level is the interplay of molecules and macromolecules. Interactions contribute to diversity and coordination of reactions to accomplish feats that would be impossible if all parts worked fully in isolation. Proteins are no exceptions and many of them undergo concerted interactions to achieve their full potential. Many interactions have been described in detail, including inter-and intra-protein domain-domain interactions, which will be the focus of this work. However, many more meaningful interactions await to be discovered and explored. An issue with the experimental description of such interactions is that many are transient and that high-throughput technologies to identify such interactions are very error prone [1]. Advances in sequencing technology and the subsequent accumulation of vast sequence databases have fueled the generation of mathematical frameworks which aim to identify protein-protein interactions [2, 3]. Some of these techniques rely on the correlated evolution of interacting proteins [4-10]. Whenever interactions are conserved across many organisms, sufficient sequence examples are now in principal available to computationally identify novel interactions relying on sequences alone.

We suggest a statistical approach based on the *coevolution of interacting protein domains*. Coevolution can be detected at very different scales, ranging from the correlated presence or absence of related proteins (or their genes) across genomes, down to the correlated usage of amino-acids in residues, which are located in different proteins but in contact across the interface. Each scale contains valuable information for detecting and understanding interactions between proteins and their domains, and adapted methods have been designed to unveil this information from data. However, none of the scales contains exhaustive information. Therefore, our work proposes a coherent mathematical-algorithmic framework bridging different scales, thereby combining the information content of the different scales.

The first, largest scale concerns the correlated presence and absence of interacting proteins in genomes. If a biological function depends on two proteins simultaneously (not necessarily via their direct physical interaction, but via any functional relation), we will either observe both proteins in a genome, i.e. the function is present, or none of them, i.e. the function is absent. More rarely we may observe the presence of only one of the two proteins. This idea is at the basis of a classical method called *phylogenetic profiling* [4, 5], which uses presence/absence correlations across genomes to predict interactions. Its accuracy suffers, however, from a number of shortcomings and confounding factors:

1. *Phylogenetic relationships* between considered genomes may introduce correlations unrelated to biological function; single evolutionary events may be statistically amplified when closely related species are included in the data. Evolutionary models taking into account the underlying species tree, have been proposed [11-13] to prune such correlations.
2. Correlations may result from direct couplings, e.g., when two domains or proteins interact physically, but they may be caused by intermediate partners: If A co-occurs with B, and B with C, also A and C will show correlations. Analyses based on partial correlations [14] and spectral analysis [15] have been proposed to *disentangle direct from indirect correlations*.
3. Simple presence/absence patterns cannot *discriminate physical interaction from more general relationships,* like co-occurrence in a biological pathway or genomic co-localization. Here, using full amino-acid sequences instead of presence/absence patterns may help to refine the analysis, e.g. via the comparison of protein-specific phylogenetic trees [6].

This last point actually suggests to change resolution, and to consider coevolution at the residue scale to refine the analysis of phylogenetic profiles. The last decade has seen important progress in this respect [16, 17], related to methods like Direct Coupling Analysis (DCA) [18, 19], Gremlin [20] or PsiCov [21]. DCA-type methods were initially developed to capture the correlated amino-acid usage of residues in physical contact. Concerning interacting proteins, they have triggered a breakthrough in using sequence covariation for inter-protein residue-residue contact prediction [16, 17], which in turn is used to guide computational quaternary structure prediction [22-25].

Beyond structure prediction, DCA was suggested for the identification of interacting proteins [9, 10, 26, 27]. Such analysis requires the construction of a large joint multiple-sequence alignment (MSA) of two protein families, with each line of the MSA containing two potentially interacting proteins. However, when proteins possess numerous paralogs inside the same genome, the matching of potentially interacting paralog pairs becomes computationally hard [8, 28]. In some cases, genomic co-localization (e.g. bacterial operons) helps to identify the interacting paralogs [18, 23, 24]. Residue-residue coevolution itself has recently been proposed as a means to match paralogs, and to identify specific interaction partners [26, 27]. While results for individual protein pairs are promising, the computational cost is prohibitive for genome-wide analysis, i.e., for systematically investigating all pairs of present protein families for signatures of coevolution and thus interaction.

Our work addresses this issue, together with Points 2 and 3 given above. We propose a common statistical-modeling framework, which is applied successively to the genomic and the residue scale (presence/absence patterns and amino-acid sequences) of coevolution. It is intended to extract information from data, which cannot be extracted at each individual scale. Performing the genome-wide analysis on the coarse scale of presence/absence patterns, we can identify promising protein-domain pairs, which are subsequently analyzed using DCA at the fine residue scale.

For the genome scale, we introduce thereby the concept of *direct phyletic couplings* into phylogenetic profiling. Using a thoroughly constructed test set of positive relations between protein domains in the bacterium *Escherichia coli*, we show that phyletic couplings substantially improve the accuracy of our prediction over mere correlations. We compare results to those obtained by a phylogeny-aware method [29], observing some interesting connections between Points 1 and 2 above. Negative phyletic couplings (i.e. one protein is typically present in a genome when the other is absent) pointedly identify alternative solutions for the same functionality, including documented cases of convergent evolution.

Still, a number of protein pairs with strong phyletic couplings are not contained in our positive test set. They constitute a first level of prediction of novel interacting partners. As discussed before, phyletic couplings are an indicator for a functional dependence between proteins. To detect direct physical interaction, we apply the DCA-based paralog matching discussed above. The resulting domain-domain pairs are highly coupled both in their joint presence across genomes, and in their evolution at the amino-acid scale. An in-depth discussion of the highest-scoring examples for such predicted but currently unknown protein-protein interactions illustrates that many of them are biologically sensible; the predictions can be tested directly in future experiments.

## Results

### Phyletic couplings improve the prediction of domain-domain relationships beyond correlations

The analysis starts with a fairly standard construction of phylogenetic profiles [5], as outlined in Fig. 1. Multiple-sequence alignments are needed at a later stage to perform inter-protein DCA. Since Pfam MSA have been extensively used in this respect, the analysis is performed on the domain level [30], using Pfam [31] as the input database. Pfam is based on reference genomes and we use the 1041 bacterial ones. The bacterial model organism *Escherichia coli* is used as a reference, i.e. only the 2682 domain families existing inside the K12 strain of *E. coli* are considered (the *Supplement* shows that the results are robust with respect to this choice). Since our method is based on covariation of presence and absence of domains in genomes, only variable domains existing in at least 5% and at most 95% of the considered genomes are considered, leaving 2041 domains. Note that the upper limit removes domains, which are omnipresent in the bacteria – mostly related to central life processes like replication, transcription and translation. However, being omnipresent, these domains cannot give any covariation signal within phylogenetic profiling. They could be analyzed using the finer residue-scale of coevolution, which might bring complementary evidence for interactions between these domains, but this analysis is out of scope in the current paper. The final input data are given by a binary phylogenetic profile matrix (PPM) of *M* = 1041 rows (species) and *N* = 2041 columns (domains), with entries 1 if a domain is present at least once in a genome, and zero if it is absent, cf. *Methods* and Fig. 1.

**Figure 1:**
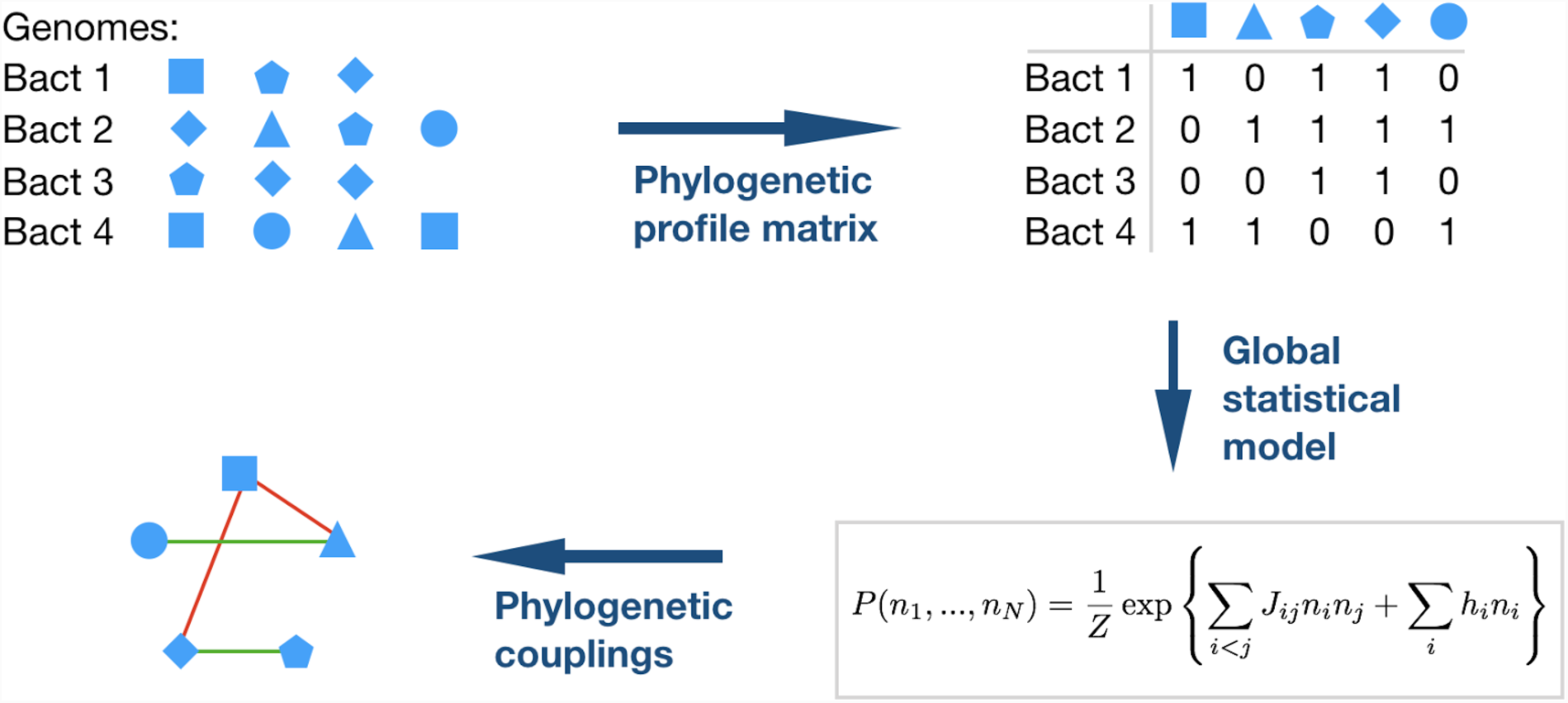
Schematic representation of the inference of phylogenetic couplings. The composition of bacterial genomes in terms of protein families is extracted from the Pfam database. The presence and absence of each family is coded into the binary phylogenetic profile matrix (PPM); note that this matrix does not account for the presence of multiple paralogs of a domain. The statistics of the PPM is reproduced by a global statistical model *P*(*n*_*1*_, *…, n*_*N*_) for a full genomic phylogenetic profile, the model corresponds to a lattice gas model in statistical physics. The strongest positive couplings (domain-domain co-occurrence) are expected to stand for positive relationships between domains, like domain-domain interactions or genomic co-localization. Negative couplings (avoided co-occurrence) is expected to indicate alternative solutions for the same biological function, like in cases of domain families in a common Pfam clan, or for convergent evolution.

An important breakthrough in coevolutionary analysis at the residue level was the step from a local correlation analysis to global maximum-entropy models [16, 32], which are able to disentangle indirect (i.e. collective) effects in correlations, and to explain them by a network of direct couplings. Here we show that the same idea can be adapted to phylogenetic profiling, and leads to a strongly increased accuracy in predicting relationships between domains. The method, which we call *Phyletic-Coupling Analysis (PhyCA)*, infers a statistical model *P*(*n*_*1*_, *…, n*_*N*_) for the phylogenetic profile of an entire species, i.e. by a binary vector (*n*_*1*_, *…, n*_*N*_) signaling the presence or absence of all *N* considered domains in the corresponding species, cf. *Methods* for details. The PhyCA model resembles a *lattice-gas model* in statistical physics, describing *N* coupled particles that can be present or absent. The phyletic coupling *J*_*ij*_ between particles / domains *i* and *j* can be positive – i.e. the presence of one domain favors the presence of the other. In this case we expect a positive relationship between the two domains, corresponding to biological processes requiring both domains. The coupling *J*_*ij*_ can also be negative – i.e. the presence of one domain favors the absence of the other. We would expect that these domains have overlapping functionalities, and one of the two is sufficient to guarantee this functionality in a species. Fig. 2A shows a histogram of the couplings found for the phylogenetic coupling matrix. We observe clear bulk of small coupling values concentrated around zero, with a broad tail for larger positive values, and a less pronounced tail for negative values.

**Figure 2:**
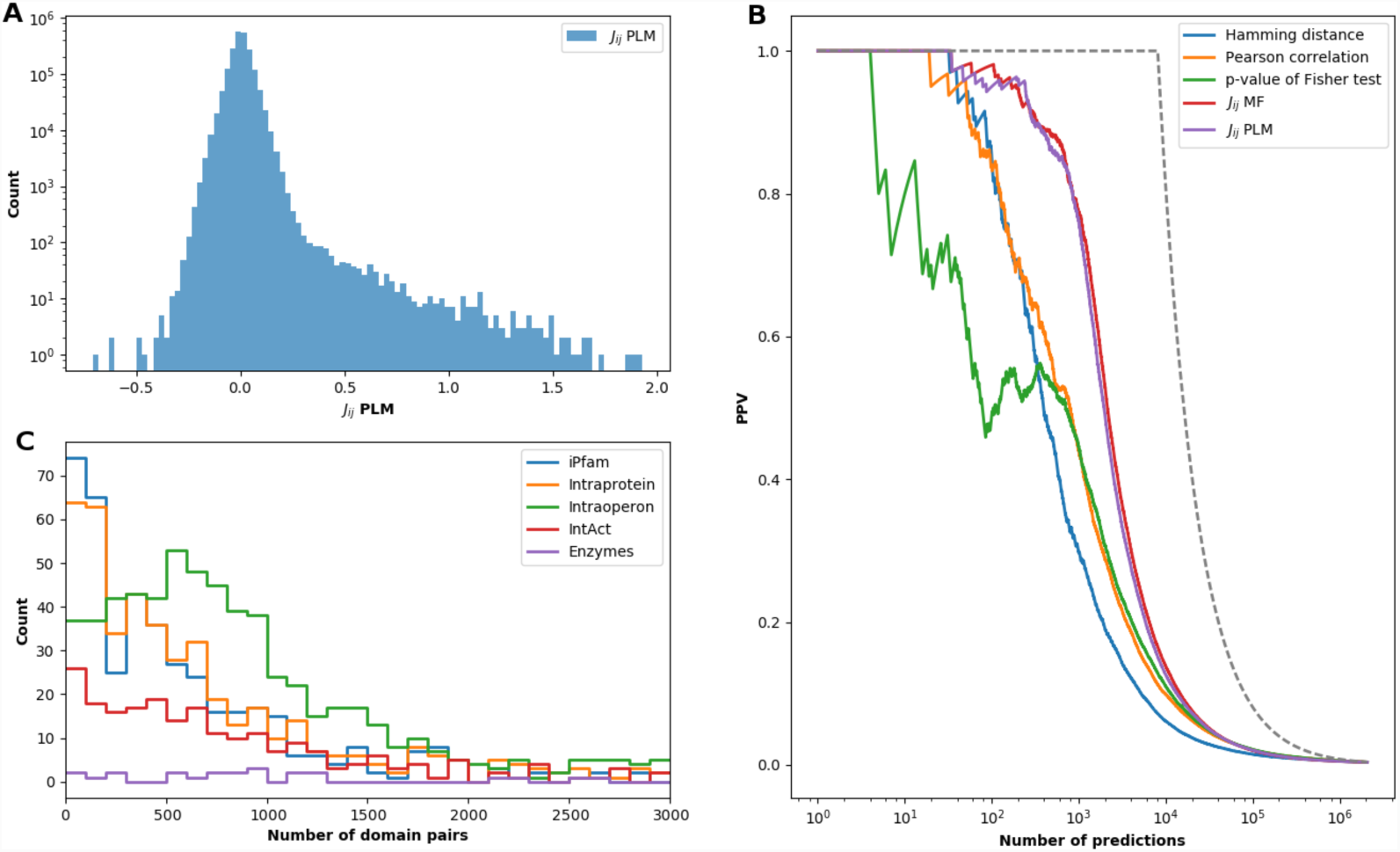
Phylogenetic couplings predict domain-domain relationships. *Panel A* shows a histogram of couplings *J*_*ij*_ as inferred using pseudo-likelihood maximization (PLM), cf. *Methods*. The histogram shows a dominant central peak around zero (note the logarithmic scale of the counts) with a pronounced fat tail for positive couplings. A small tail for negative couplings is visible, too, but much less pronounced. *Panel B* shows the PPV (positive predictive value), defined as the fraction of known domain-domain relations in between the strongest couplings or correlations. A random prediction would correspond to a flat line close to zero; a perfect prediction would follow the dashed black line. Note that the curves corresponding to phylogenetic couplings (inference vis PLM or MF (mean field), cf. *Methods*) are substantially higher than those using correlation measures. *Panel C* shows, in bins of 100 domain pairs ordered by their phyletic couplings, the number of pairs belonging to the different parts of the positive-relation list (note that the categories are not exclusive, so the sum of different categories may exceed 100). We find enrichment of co-localized and interacting domain pairs, but not of related enzymes.

The performance of PhyCA can be assessed by comparing the domain pairs of strongest phyletic couplings to a carefully compiled list of 8,091 known domain-domain relations. As is explained in *Methods*, we have included genomic, functional and structural relationships: Domains may coexist inside a single protein, they may be co-localized in an operon, they may be in contact in an experimental crystallographic structure or an interaction might be known according to other experimental techniques, or they may belong to enzymes catalyzing related reactions.

The PhyCA couplings *J*_*ij*_ are ordered by size, and the fraction of positive relations in between the highest-scoring domain-domain pairs is calculated (PPV = positive predictive value). Fig. 2B shows the results: we observe a strong enrichment in known positive relations in between strongly phyletically coupled domain-domain pairs. This enrichment is much stronger than for local correlation measures like Hamming distance, Pearson correlation or p-value of Fisher’s exact test applied individually to two domains (i.e. two columns of the PPM): E.g., for the first 1000 predictions we observed a PPV of about 0.8 for the phyletic couplings, and only 0.3-0.5 for the different correlation measures. As is shown in Fig. 2C, interacting and co-localized domain pairs are enriched in the predictions of large positive couplings, whereas enzymes from related metabolic reactions are not.

Databases of genome-wide protein-protein or domain-domain interactions are currently incomplete. We therefore expect the real PPV to be even higher than the one measured in Fig. 2: strongly coupled domain-domain pairs *not* belonging to our list of positives may actually be considered as predictions for new, currently unknown relations. According to the observations in Fig. 2C, these relations might be direct physical interactions, but also genomic co-localization (frequently related to joint biological function). Before exploring these possibilities in more detail and on the finer scale of the residue-residue coevolution, we compare the PhyCA results to phylogeny-aware correlation analysis and investigate the negative tail of the *J*_*ij*_ distribution.

### Comparison of phyletic couplings to phylogeny-aware analysis of correlated presence/absence patterns

Phyletic couplings are, like simpler correlation measures, based on counting co-presence and co-absence of proteins or domains. However, due to the uneven phylogenetic distribution of species in our dataset, single evolutionary event may be amplified when appearing in an ancestor of several closely related species. More importantly in the context of this study, phylogeny may introduce spurious correlations in the presence and absence of domains, which are not related to biological function.

To remove this bias, several methods have been proposed, cf. [11, 13], which use evolutionary models to decide, if observed correlations can be explained by phylogeny alone (i.e. by independent evolution on a phylogenetic tree), or remain significant even when such phylogenetic effects are removed. Since this idea is complementary to the one behind PhyCA, it is important to compare the outcome of both approaches.

To this end, we have used the CoPAP (coevolution of presence-absence patterns) server [29]. It uses the same type of binary input matrix of our approach, and is able to efficiently treat matrices of more than 2,000 domains across more than 1,000 species. As an output, CoPAP provides p-values measuring the significance of correlated domain presence and absence, as compared to independently evolving domains on the same phylogenetic tree. The group of maximum significance (*log*_10_*p* < –7.9) contains 3,611 domain pairs, out of which 1,251 (34.6%) are true positives in our list of known domain-domain relationships.

Since a further sorting of these pairs using CoPAP results is not possible (p-values are calculated using finite simulations), we compare them to the first 3,611 domain pairs extracted by PhyCA, and to the 3,611 domain pairs of highest Pearson correlation. The Venn diagram in Fig. 3 allows for a number of interesting observations:

**Figure 3:**
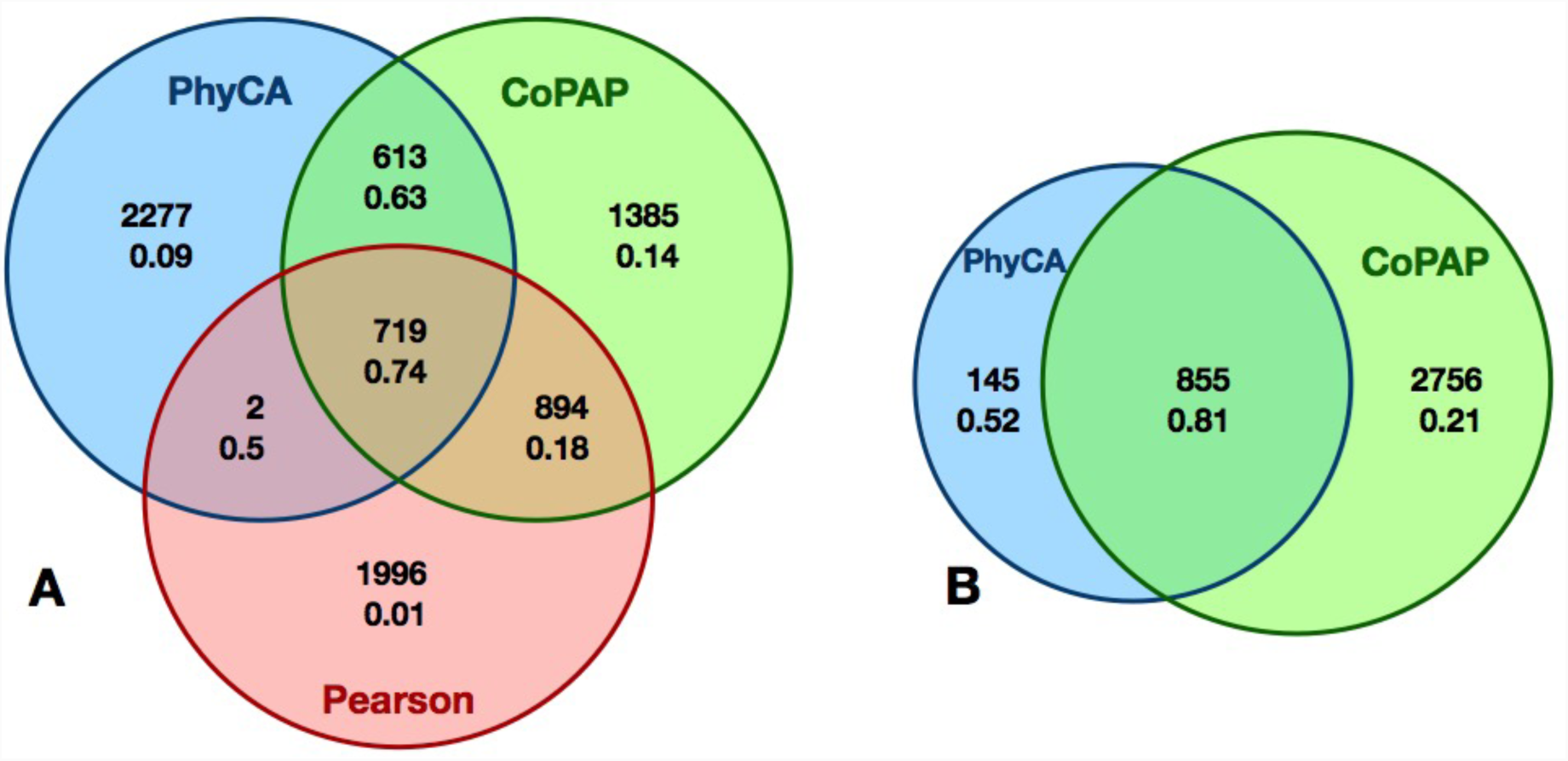
Comparison of simple correlations, phyletic couplings and phylogeny-corrected correlations. Panel A shows a Venn diagram for the 3,611 first predictions of each of the three coevolution measures as extracted by Pearson correlation (red), PhyCA (blue) and CoPAP (green). Numbers are the size of the corresponding intersection, and the PPV indicating the fraction of true positives according to our list of positive domain-domain interactions. Panel B compares the first 3,611 CoPAP predictions of highest possible significance, with the most significant 1,000 PhyCA predictions. Most of them (855) are found to be significant by CoPAP, and of very high PPV (81%). However, not all CoPAP pairs are strongly coupled, and thus PPV is reduced (21%).

- While CoPAP and PhyCA have similar global PPV, with an advantage for CoPAP (34.6%) over PhyCA (31.2%), Pearson correlation performs substantially worse (PPV 19.7%).
- Very small fractions of the correlated pairs, which are discarded by PhyCA or CoPAP, are TP: PhyCA discards 2,890 pairs of PPV 6%; CoPAP discards only 1,998 pairs, but with even lower PPV (1.2%).
- 74% of the 721 correlated pairs, which are retained by PhyCA, are TP. Note that almost all of them (719/721) show also a significant CoPAP signal.
- Only 43% of the correlated pairs, which are retained by CoPAP, are TP. PhyCA divides them into two groups of comparable size but distinct PPV. For the 719 pairs retained also by PhyCA, the PPV rises to 74%. The other 894 pairs have weak phyletic couplings, so their significant correlation has to be interpreted as dominated by indirect effects. Actually only 18% are TP.
- When going to lower Pearson correlations, both CoPAP and PhyCA decrease their accuracy. However, their intersection shows 613 pairs with a high PPV of 63%.
- The 2,277 pairs only identified by PhyCA have a low PPV of only 9%. This is coherent with Fig. 2B, which shows a sharp PPV drop in PhyCA after the first ca. 1,000 phyletic couplings. We have therefore compared these 1,000 domain pairs separately to CoPAP. A vast majority of 855 pairs have the highest possible significance in CoPAP, this intersection has a PPV of 81%. The other 15% have lower CoPAP scores and lower PPV (52%). Interestingly, only 21% of the 2,756 strongest CoPAP without strong coupling are TP, illustrating again the capacity of PhyCA to – at least partially – disentangle direct couplings from indirect correlations.

In principle, CoPAP and PhyCA treat very different confounding factors of coevolutionary analysis – phylogenetic biases and indirect correlations. So, it might appear astonishing that almost none of the correlated pairs, which are strongly coupled in PhyCA, are actually discarded by CoPAP. The reason might be given by the spectral properties of the covariance matrices of the input data, and their relation to phylogeny and direct couplings. As shown in [33], the phylogenetic bias is most evident in the largest eigenvalues of the data-covariance matrix. These correspond mostly to extended eigenmodes, which in turn give rise to a dense network of small couplings [15, 34]. On the contrary, the strongest pairwise couplings are related to small eigenvalues with more localized eigenmodes, which give rise to strong, sparse couplings. Phylogenetic biases and strong direct couplings are thus related to different tails of the eigenvalue spectrum of the covariance matrix, the strongest PhyCA couplings are thus robust with respect to phylogenetic biases. On the other hand, there are non-phylogenetic but indirect correlations, therefore PhyCA separates the CoPAP output into strongly coupled pairs of high PPV, and weakly coupled pairs of reduced PPV.

### Negative phylogenetic couplings appear between alternative solutions for the same biological function, including cases of convergent evolution

A smaller tail of negative phylogenetic couplings can be observed in Fig. 2A. A negative coupling disfavors the joint presence of two domains in the same genome, i.e., if one of the negatively coupled domains is present in a genome, the other is less likely to be present, too. Intuitively this suggests similar functionalities, one of the two domains is sufficient, the joint presence unnecessary or even costly for a bacterium. Such pairs, called anti-correlogs in [14] were used in [35] to identify analogous enzymes replacing missing homologs in biochemical pathways.

When using *E. coli* as a reference genome, the number of such negative couplings is limited, since only domain pairs co-occurring in *E. coli* are analyzed. To better understand the meaning of negative couplings, we have therefore extended the original analysis to all 9,358 families containing bacterial protein domains. While results restricted a posteriori to *E. coli* are very robust (96% correlation, cf. *Supplement*), the extended analysis leads to a substantially higher number of negative couplings.

To explore these in some detail, we analyzed the 20 domain pairs with the strongest negative couplings, cf. Table 1 (an extended list is given in the *Supplement*). From their detailed analysis it is evident that protein pairs can be classified into three distinct groups. First, we find several cases of convergent evolution as evidenced by proteins with the same or similar activities but distinct protein structures (rankings 1, 2, 9, 14, 15, 16). Second, we find domain pairs of the same fold and, where known, of similar activity. For various reasons these are not described by the same Pfam HMM (rankings 3, 4, 6, 7,8, 10,11, 17,19), but typically belong to the same Pfam clan indicating distant homology. Lastly, there are several cases of relatively unknown activity, and some domains have no known structure (rankings 5, 12, 13, 18, 20).

**Table 1:**
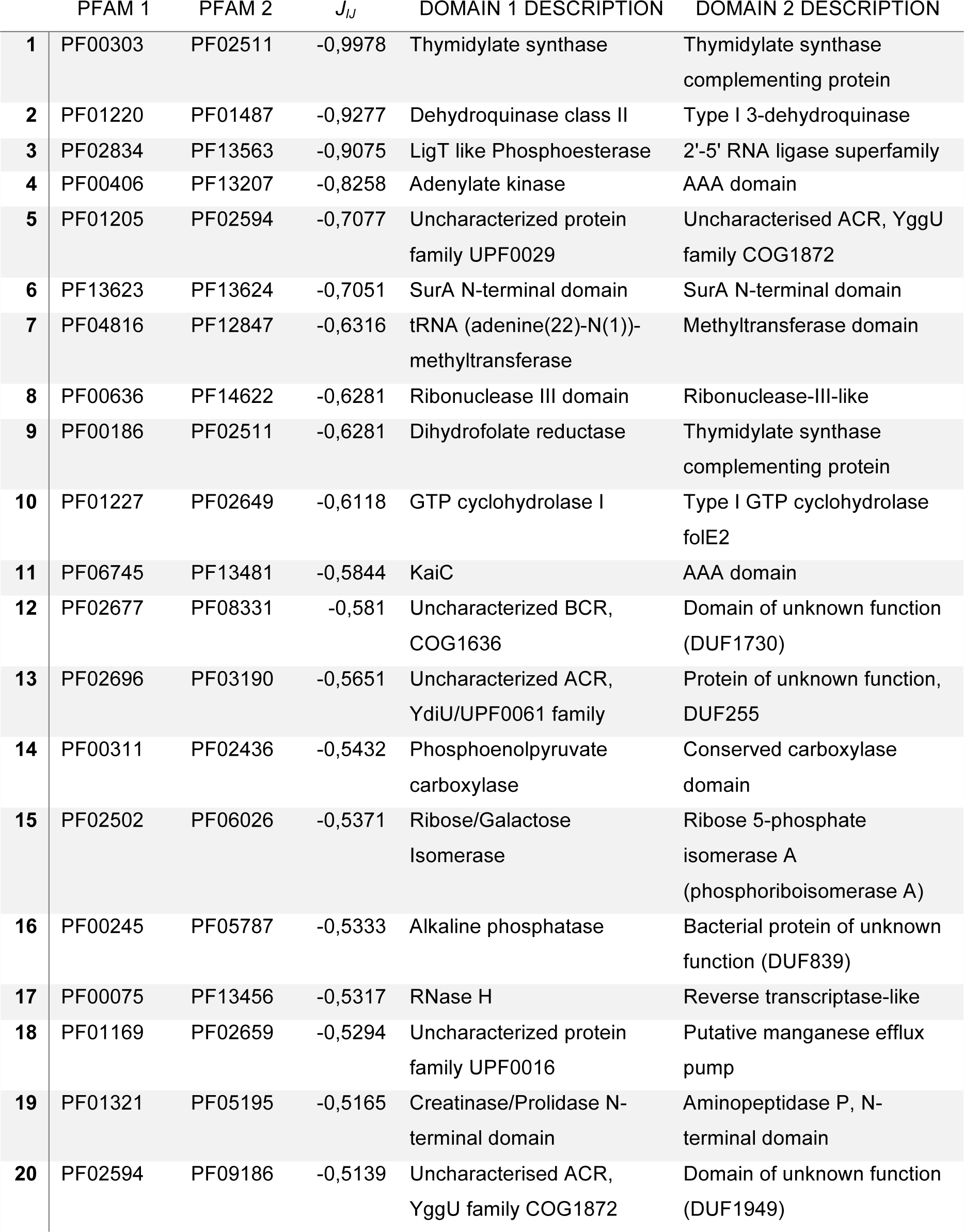
The 20 domain pairs of top negative phyletic couplings.

Cases of convergent evolution include PF00303 and PF02511, which describe two different thymidylate synthases, the former a 5,10-methylenetetrahydrofolate, the latter a flavin dependent enzyme [36]. Interestingly, PF00186, dihydrofolate reductase is also strongly negatively coupled with PF02511 (but positively to PF00303), since the former is not needed to regenerate 5,10-methylenetetrahydrofolate when the flavin-dependent enzyme is used. Other cases of convergent evolution are PF01220 and PF01487 that describe two classes of dehydroquinases with similar activity but significantly different primary and secondary structure [37]. PF00311 and PF02436 describe proteins in oxaloacetate biogenesis, the former from phosphoenolpyruvate, the later from pyruvate and ATP. PF00245 and PF05787 describe two classes of bacterial alkaline phosphatases, termed PhoA and PhoX with distinct protein folds [38]. PF02502 and PF02436 distinguish two classes of ribose-or phosphoribo-isomerases with differing enzyme folds.

Structurally similar proteins that are identified by different Pfam families are of less interest and will not be separately described. The fact that they are distinct enough in sequence to be covered by separate Pfam families suggests a level of divergent evolution, i.e. one or the other domain has distinct features such us additional interaction partner, distinct activity regulation etc.

Of special interest are domain pairs with unknown function. Ideally, if the function of one Pfam family becomes available one can infer the function of the other family as well. In addition, the evolutionary importance of a given protein family and its activity is often judged by its conservation across different phyla and organisms. This however neglects cases of unknown convergent evolution. Among the highest negatively coupled pairs, we did not find any, where the function of one has been clearly identified and the function of the other has not. However, there are several instances, where a potential role has been loosely associated with one or the other domain. For instance, PF01205 and PF09186 have been suggested to be involved in countering translation inhibition under starvation conditions [39]. These domains are strongly negatively coupled with PF02594, suggesting that the latter might also serve a role in countering translation inhibition. PF01169 and PF02659 are both putative transporters, the former for calcium [40], the latter for manganese ions [41]. Their coupling suggests overlapping specificities or roles. PF02677 and PF08331 describe two entirely unstudied bacterial proteins. The later appears associated with iron-sulfur cluster domains, suggesting a potential role in redox regulation. Lastly, we find a negative coupling between domains PF02696 and PF03190. Both proteins are entirely unstudied in bacteria, but they are also common in Eukaryotes where the latter is a proposed redox protein that has been implicated in fertility regulation in mammals [42]. It would be interesting to unveil their function in the bacteria.

In summary, where known, negative couplings are associated with function and potential new function might be derived from this list.

### A residue-scale DCA analysis of phylogenetically coupled domain pairs unveils directly coevolving pairs

As seen in Fig. 2C, a large positive phyletic coupling is a strong signal for a positive relationship between two domains, but not necessarily for a direct physical interaction of the two domains within a protein complex. Furthermore, co-localization of two domains either inside the same protein (i.e. an evolutionary conserved protein architecture) or inside the same operon may lead to strong phyletic couplings.

Relying only on the coarse scale of coupled presence and absence in genomes, does not reveal more detailed information. Since the number of domain-domain pairs under question is limited as compared to all domain pairs existing in *E. coli*, we can afford computationally more expensive approaches, which study coevolution of domain pairs at the individual residue scale. To this effect, we use the procedure suggested in Gueudré et al. [27]: Two Pfam MSA for the two domain families are matched using a variant of DCA such that (a) only sequences appearing inside the same species are matched and (b) the inter-domain covariation as measurable by DCA is maximized. In [27] it was shown that this idea allows to identify protein-protein interactions via a large coevolutionary score between the two domains at a sufficiently large joint MSA. DCA scores above 0.2 at an effective sequence-pair number of at least 200 (sequences below 80% sequence identity, cf. *Supplement*) can be considered as a strong indicator for a potential interaction [10, 27]. On the contrary, according to [43], a low DCA score is not necessarily a sign for the absence of a physical interaction. A low score might also originate from a relatively small or structurally not well conserved interface, both resulting in a weak coevolutionary signal.

We have applied the progressive paralog matching procedure of [27] to the 500 most strongly coupled domain pairs, which are not in our previously constructed test set of positive domain-domain relations, i.e. to the first 500 *prediction*s at the scale of phyletic couplings. The results are presented in Fig. 4A: 360 domain pairs have an *M*_*eff*_ above 200, and DCA results can thus be considered reliable. Of those 45 pairs have an inter-domain DCA score above 0.2 (24 put of the first 200 PhyCA predictions). This number is significantly larger than randomly selected protein pairs, cf. Fig. 4B: only 10 pairs have a score above 0.2 and *M*_*eff*_ above 200, mostly related to short amino-acid sequences. This shows that the preselection by high phylogenetic couplings leads to a subsequent enrichment of high DCA scores also at the residue scale. For comparison, we have also applied the matching procedure to the 200 domain-domain pairs, which are known to interact by iPfam [44], and which have high phylogenetic couplings, cf. Fig 4C. 29 have a significant DCA score at large enough sequence number. Interestingly, the signal is only marginally stronger than for the newly predicted relations, which are discussed in more detail below.

**Figure 4:**
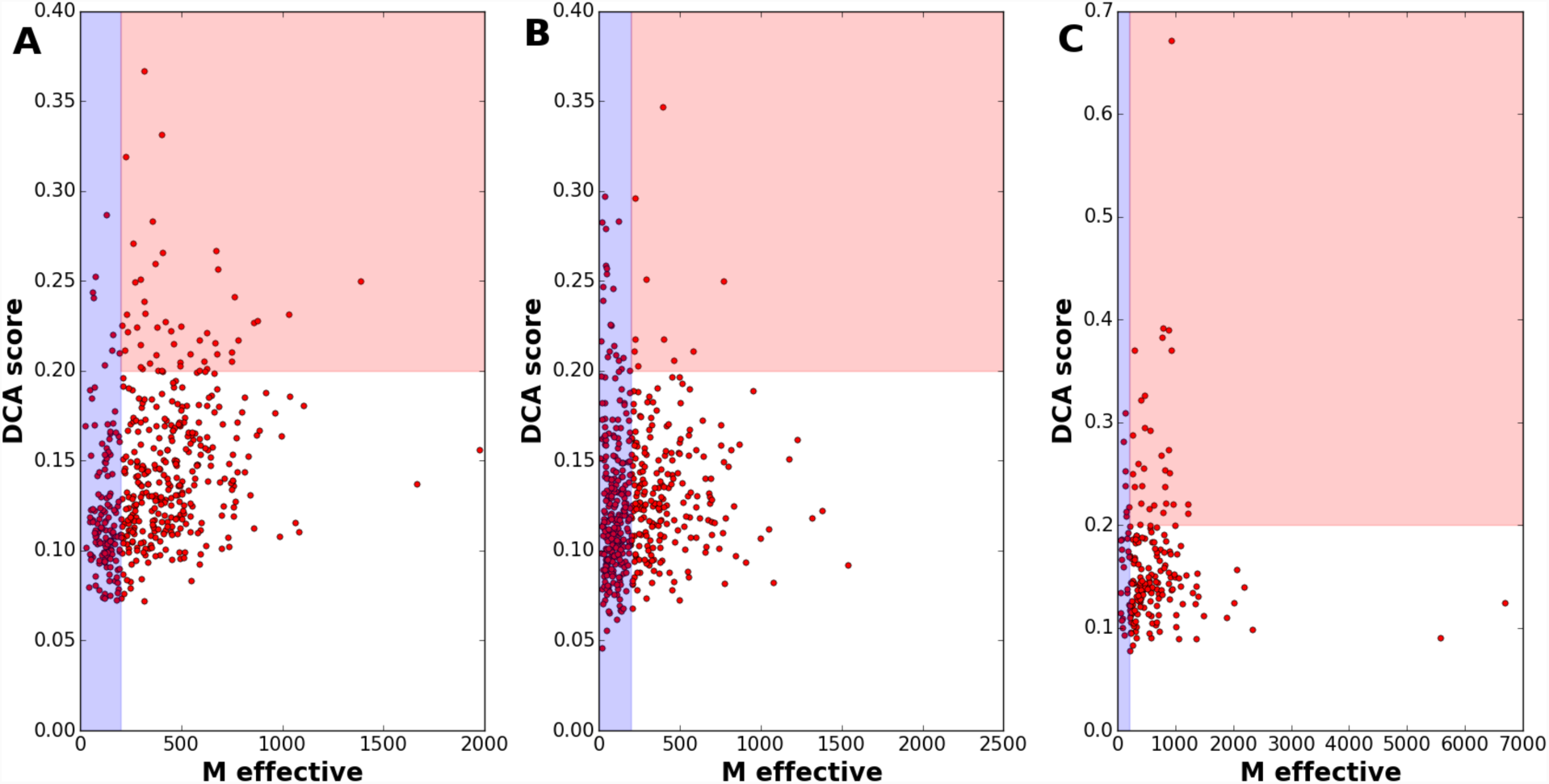
DCA identifies strong residue-scale coevolution between phyletically coupled domain pairs. Panel A shows the effective sequence number (defined as the sequence number at 80% maximum sequence identity, cf. *Supplement* for the precise definition) and the DCA scores for the 500 domain pairs of strongest phyletic coupling not belonging to the positive-relation set (i.e. the 500 most significant predictions). The interesting region is the red one, where sequence numbers are sufficient to provide reliable DCA results, and DCA scores are beyond 0.2 as established in [10]. Panel B shows, as a comparison, the results for 500 randomly selected domain pairs. Only very few pairs show substantial scores, most of them related to very short peptides. Panel C shows a positive control, the 200 pairs of highest phylogenetic couplings belonging to iPfam are analyzed analogously. The fraction and the amplitude of high DCA scores is slightly increased with respect to Panel A, but the qualitative behavior is similar.

## Discussion

In this work, we propose a coevolutionary analysis connecting signals at the phylogenetic level (correlated presence of domain pairs across genomes) with the residue level (correlated occurrence of amino acids between proteins). At the phylogenetic level, we introduce the concept of *phyletic couplings*: By using a global statistical model, we are able to disentangle direct and indirect correlations in the presence and absence of protein domains across more than 1000 fully sequenced representative bacterial species. Couplings substantially increase the capacity to find relations between domains beyond correlations; these relations can be physical interactions, but also genomic co-localization (and thus likely functional relations). Standard correlation measures used in phylogenetic profiling only reach 30-50% of true positives between the first 1000 predictions. In contrast the positive predictive value of phylogenetic couplings reaches about 80%. The results are very robust: when applying the same methodology to all 9358 Pfam domains appearing in the bacteria, and selecting only later the couplings between domains present in *E. coli*, couplings have 96% correlation with the couplings found by the procedure described before.

The high accuracy of phyletic couplings in predicting domain-domain relations, along with the robustness of these couplings when extensively changing the data set, allows us to hypothesize that large couplings not corresponding to known relations predict novel, unknown relations. A list of the 500 first predictions is provided in the *Supplement*.

As mentioned, a large phyletic coupling does not automatically imply a direct physical interaction. Two proteins may have a strong phyletic coupling because they belong to the same multi-protein complex, without touching each other. They may have a strong phyletic coupling, because they act within the same biological process or pathway, again without any direct interaction. To refine the results and predict physical interactions, we have added a coevolutionary analysis on the scale of residue-residue covariation, as provided by DCA, in the version with paralog matching as recently proposed in [27]. We find that 72% of the 500 phylogenetically most coupled pairs correspond to large enough alignments to run DCA, and 12.5% of these have significant DCA scores.

These domain pairs are our strongest candidates for predicted domain-domain interactions. Since they are not co-localized in the same protein, they also provide predictions for new protein-protein interactions. We analyzed in detail the 24 pairs with a score larger than 0.2, which result from the first 200 PhyCA predictions filtered with DCA.

Among these 24 pairs we find several examples of known interactions that have not yet been structurally resolved. These include K^+^ transporter subunits KdpC (PF02669) and KdpA (PF03814) [45], Sigma54 activator (PF00158) and Sigma54 activator interacting domain (PF00309) [46] and exonuclease VII subunits domains PF02609, PF2601 and PF13742 [47].

For several additional positively coupled pairs an interaction seems functionally very likely but to our knowledge no interaction studies are available. These are all proteins involved in pilus formation or maturation. Domain PF06750 is a putative methyl transferase domain in the prepilin peptidase PppA, and proposed to interact with methylation motif domain PF07963, found in numerous pilin proteins and with PF05157, a type II secretion system protein [48, 49]. PF05157 is also predicted to interact with domain PF05137 found in the PilN fimbrial assembly protein required for mating in liquid culture [50].

Of interest, there are predicted interactions for several members of biosynthetic pathways catalyzing either consecutive or closely following reactions. These include domains PF02542 and PF13288 of isoprenoid biosynthesis enzymes Dxr and IspF, domains PF00885 and PF00926 of riboflavin biosynthesis enzymes RisB and RibB and domains PF01227 and PF01288 of tetrahydrofolate biosynthesis enzymes Gch1 and HppK. A more complex connection is predicted between multiple domains of molybdenum cofactor biosynthesis enzyme MoaC (PF01967), MoeA (PF03453 and PF03454) and MoaA (PF06463). Similarly, scores suggest a protein-protein interaction between domains of hydrogenase maturation enzymes HypF (PF07503) with HybG (PF01455) and HycI (PF01750).

Perhaps most intriguing are the observation of strongly coupled co-occurrence and potential protein-protein interactions of two proteins pairs. Ada (PF02805) and AlkA (PF06029) are two enzymes involved in DNA repair in response to alkylation damage [51, 52]. One of the proteins serves as demethylase of guanosyl residues whereas the other excises alkylated nucleotides. These seemingly complementary functions suggests that an interaction is plausible. The other pair is YoeB (PF06769) with HicA (PF07927). These two proteins constitute two toxins of distinct toxin-antitoxin systems. Both proteins inhibit translation by distinct and complementary mechanisms and an interaction seems plausible. YoeB blocks the ribosome A site leading to mRNA cleavage [53]. HicA interacts with mRNA directly and thus acts independent of the translation apparatus [54].

Additional and perhaps plausible interactions are predicted between domains PF05930 and PF13356 of prophage protein AlpA and several phage integrase proteins as well as between domain PF13518 with PF13817, the former a HTH domain commonly associated with transposase domains and the latter a transposase domain.

Insufficient information on the function of two domain pairs and their associated proteins does not allow us to draw any conclusions on the plausibility of interaction. These are for domains PF02021 and PF13335 of proteins YraN and YifB and domains PF01906 with PF02796, the former a metal binding domain and the latter a domain found in site specific recombinases.

Lastly, we find three proposed interactions between domains found in ribosomal proteins RL36, RL34 and RL32 (PF00444, PF00468, PF01783) and also a protein of unknown function YidD (PF01809). We consider these to be likely false positive predictions since we previously observed spurious results for members of very large macromolecular complexes such as the ribosome [10]. At least the interaction between YidD and RL36 seems plausible, as the former has been suggested to play a role as membrane protein insertion factor [55].

In summary, we are able to recapitulate several known or plausible but structurally unresolved interactions and find several examples of interaction that should be of interest for future experimental studies.

Similarly, negative phylogenetic couplings appear to be biologically reasonable. They disfavor the joint presence of two domains within the same genome. In our analysis of the pairs of the strongest negative couplings, presented above in *Results*, we actually find many pairs having the same functionality, including documented pairs of convergent evolution. Some pairs actually are of unknown function, and our method might help to transfer functional annotations from one domain to the other.

An important extension would be the application of our approach beyond the bacteria. Bacteria, due to their compact genomes, are overrepresented in genomic databases, including the Pfam database, which we used for our analysis. To test the applicability to higher organisms, we have repeated the same procedure, concentrating on eukaryotic genomes and taking humans as the reference species. Data get much less abundant; the phylogenetic profile matrix now contains 5343 domains as compared to only 481 eukaryotic species. Still, phyletic couplings, when compared to a positive list extracted from domain architectures of human proteins (co-localization in one protein), from iPfam [44] and human entries in IntAct [56], show a similar performance as the bacterial case, cf. Fig. 5A: 75% of the first 1000 couplings correspond to known domain-domain relations. Entries corresponding to protein-protein interactions (iPfam, IntAct) are again significantly enriched, even if to a lesser extent than in the bacterial case. The most important difference emerges, however, when using paralog matching and DCA on the 200 most coupled predictions (i.e. pairs with strong phylogenetic coupling but not belonging to the positive list), cf. Fig. 5B: Only 2-4 have sequence numbers that allow for reliable DCA results. More eukaryotic genomes are urgently needed to carry out our full procedure also in higher species.

**Figure 5:**
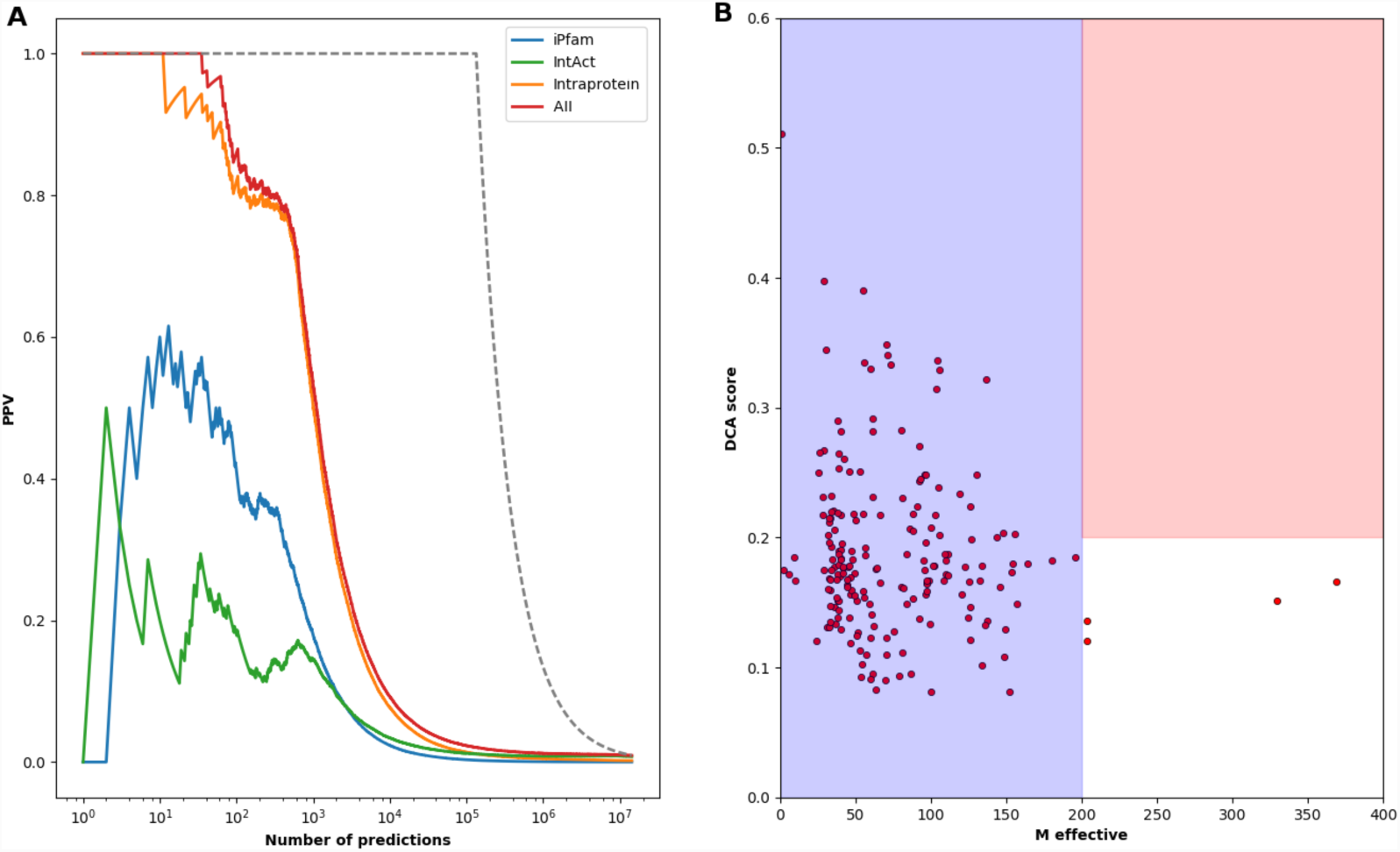
Performance of our multi-scale coevolutionary analysis for human protein domains. Panel A shows the positive predictive value of the phyletic couplings for predicting positive domain-domain relationships (including protein architecture, iPfam and human IntAct entries). While there is a clear overrepresentation of intra-protein localization in between the highest-scoring domain pairs, also physical interactions as captured by iPfam and IntAct are enriched in particular in the first ca. 10^3^ phyletic couplings. The overall performance is coherent with the one found in the bacteria. Panel B shows the paralog-matching and DCA results for the 200 most coupled domain pairs, which are not in the positive-relation dataset. We observe that currently the joint MSA are too small (Meff < 200) to allow for a reliable application of DCA to detect inter-protein residue-scale coevolution.

To conclude, our work illustrates the potential of combining rapidly growing genomic databases and statistical modelling: the increasing number of fully sequenced genomes allows for extracting rich *samples for the variability in protein content and protein seque*nces across hundreds and thousands of species; their statistical analysis helps us to detect multiple scales of coevolution between interacting or functionally related proteins.

The *genomic scale* explores the correlated presence or absence of proteins (in the sense of homologous protein families) across species. This correlation has been used before within phylogenetic profiling to detect functional relations or direct interactions between proteins. Within our work, we propose to infer direct phyletic couplings via global statistical models, and prove that this concept strongly improves our capacity to detect protein relations over local correlation measures.

However, phylogenetic couplings cannot distinguish between functional relations or direct interactions between proteins. This problem can – at least partially – be resolved at the *residue scale* of inter-protein coevolution. Interacting proteins show a correlated usage of amino acids across their interface, and again global statistical modelling approaches like DCA have been used to discriminate between interacting and non-interacting protein pairs.

Since the computational cost of the residue-residue scale analysis is high, it is possible to analyze all pairs of the order of 10-50 proteins, but not all pairs of the order of thousands of proteins forming a species’ proteome. It is the combination of both scales, which allows us to first explore the genomic scale and then refine promising results at the residue scale. Doing so, we have provided a number of biologically sensible predictions for currently unknown protein-protein interactions. We provide a list of these predictions, which in turn may be tested directly.

## Methods

### Phylogenetic profiles

Data are extracted from the Pfam 30.0 database [31]. For each of the 1,041 bacterial genomes present in Pfam, we extract all appearing protein-domain families, accounting to a total of 9,358 Pfam families. A restriction to *Escherichia coli* as reference genome (i.e. counting only domains contained in *E. coli*) reduces this to 2682 domain families. Since we are interested in the *correlated* presence / absence of domains across species, we remove all domain families with less than 5% or more than 95%, keeping only domains with at least 53 and at most 988 appearances. This removes in particular omnipresent domains related, e.g., to replication, transcription and translation. The final phylogenetic profile matrix (PPM) is a binary matrix containing *M* = 1,041 bacteria and *N* = 2,041 domains. Entries are one if a domain is present in a species (at least once), and zero if it is absent. Note that a zero entry typically indicates a true absence of the domain in a genome, since the profile matrix is entirely built on fully sequenced genomes.

In standard phylogenetic profiling [5], correlations between domains are evaluated via the Hamming distance, Pearson correlation or p-values of Fisher’s exact test, cf. the *Supplement* for the definitions in the context of our work.

### Phyletic couplings

In analogy to the direct-coupling analysis on the level of amino-acid sequences, we model the phylogenetic profiles via the maximum-entropy principle by a global statistical model

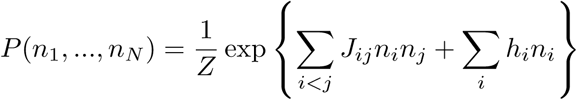

with (*n*_*1*_, *…, n*_*N*_) being a binary vector characterizing the presence (*n*_*i*_ = 1) or absence (*n*_*i*_ = 0) of domain *i* in a species, and *Z* is a normalization constant also known as partition function in statistical physics. The *phyletic couplings J*_*ij*_ and *biases h*_*i*_ are to be determined such that the model *P* reproduces the one-and two-column statistics of the PPM 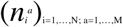:

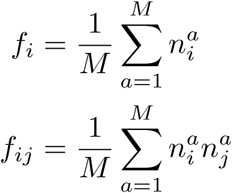

with *f*_*i*_ being the fraction of genomes in the PPM carrying domain *i,* and *f*_*ij*_ the fraction of genomes containing both domains *i* and *j* simultaneously. While the exact determination of the marginal distributions of P requires exponential-time computations, we apply the mean-field (MF) and pseudo-likelihood maximization (PLM) approximations successfully used in the context of DCA [19, 57]; cf. the *Supplement* for technical details. Strong positive couplings favor the joint presence or joint absence of two domains, signaling therefore a positive association between the two (genomic colocalization, functional relation, domain-domain interaction). Strong negative couplings favor the appearance of only one out of the two domains, signaling domains of similar function (e.g. convergent evolution).

Before analyzing the phyletic couplings, we apply the so-called Average Product Correction (APC) [58], cf. *Supplement*. APC is widely used to suppress spurious couplings resulting from the heterogenous conservation statistics domain families across genomes (cf. [59]) as compared to functional couplings.

### Direct coupling analysis of inter-protein residue coevolution

To assess the coevolution on the finer scale of residue-residue coevolution, we have applied exactly the progressive matching and analysis procedure recently published by part of us in [27], details about the procedure are given in the *Supplement.* It starts with two domain alignments, containing only bacterial protein sequences. It matches sequences between the domain families, such that (a) only sequences from the same species are matched and (b) the total inter-family covariation signal is maximized. Results are considered positive if (i) the effective number of matched sequences (at 80% seq ID) exceeds 200 and (ii) the covariation score exceeds 0.2. It has been established in [10, 27] that larger scores are rarely obtained by unrelated protein families. Note that a smaller score may be related to a functional relationship rather than a physical protein-protein interaction, or also to a small or non-conserved interaction interface [43].

### Known domain-domain relationships

To assess the accuracy of our predictions, we have compiled a number of known relationships (provided in *Supplement*). They come from different databases, the same domain-domain pair may appear multiple times, but it is counted only once in the final list of positives:

1. *Intra-protein localization*: From the Pfam database [31], we have extracted a list of domain pairs, which co-occur inside single proteins in *E. coli*. Out of 3,116 proteins, 952 contained multiple domains, giving rise to 799 distinct domain-domain relations.
2. *Intra-operon localization:* Proteins, which are co-localized inside operons, frequently share at least part of their biological function. Using a list of operons from *E. coli* [60], we compiled a list of 4,087 colocalized domain pairs.
3. *Protein-protein interaction:* The IntAct database [56] contains 5,318 pairs of experimentally found protein-protein interactions. At the domain level, we pair all domains in one protein with all domains in the second protein (adding possibly unrelated domain pairs to those interacting), obtaining 3,070 domain pairs.
4. *Domain-domain contacts in 3D structures:* The iPfam database [44] contains domain-domain interactions extracted from structural domain-domain contacts in experimentally determined complex structures in the PDB. We included intra-and inter-chain contacts, i.e. domain-domain contacts inside a protein or between two proteins. Note that this list does not refer to *E. coli* as reference genome. In total, this accounts to 545 know relationships.
5. *Metabolic relationships between enzymes:* Using the reconstruction iJR904 of *E. coli’s* metabolic network [61] and filtering out “currency” metabolites involved in more than 50 reactions (such as water, ATP etc.), we considered three relationships:

a. *common substrate –* pairs of enzymes catalyzing reactions with at least one common substrate;
b. *common product –* pairs of enzymes catalyzing reactions with at least one common product;
c. *reaction chains –* pairs of enzymes catalyzing subsequent reactions, i.e., one product of one reaction is substrate of the second.

This lead to a total of 677 known relationships.

The total list contains 8,091 domain-domain pairs, as compared to the 2,081,820 possible pairs, which can be formed out of the 2,041 domains in our PPM.

## Availability of data and materials

A *Supplement* containing supplementary method information and results is available via the journal’s web page. The code for estimating phylogenetic couplings and data for results (list of positive domain-domain relations, phyletic couplings for bacteria with and without *E. coli* as reference, for eukaryotes with human reference, DCA-scores for top 500 new predictions by phylogenetic couplings) are provided in the GitHub repository https://github.com/GiancarloCroce.

